# Who said that? Deciphering Complex Coral Reef Soundscapes with Spatial Audio and 360° Video

**DOI:** 10.1101/2024.12.16.628659

**Authors:** Marc S. Dantzker, Matthew T. Duggan, Erika Berlik, Symeon Delikaris-Manias, Vasileios Bountourakis, Ville Pulkki, Aaron N. Rice

## Abstract

Coral reef soundscapes hold an untapped wealth of biodiversity information. While they are easy to record, they are nearly impossible to decipher because we know very little about which sound is made by which species. With identified fish sounds, acoustic monitoring can directly measure biodiversity, detect key or invasive species, identify behavioral events and frequencies, and estimate abundance, all at temporal and spatial scales not possible with methods like eDNA or visual surveys. Using a novel approach to visualize underwater sound, we present the most extensive collection of identified natural sounds from reef-associated Atlantic fishes. These sounds were all ascribed to species *in situ* on a crowded Caribbean coral reef. The soundfield analysis technique combines visualizations of spatial audio with 360° video recordings, a method not previously accomplished underwater. We used these species identifications to decipher a representative section of a soundscape recording from a separate recording device. We have collected our identified recordings into a growing open-access resource to catalyze passive acoustic monitoring research, enabling a species-specific resolution of coral reef soundscape dynamics and providing critical validated information for developing machine learning models required to analyze an ever-expanding collection of long-term recordings.

## Introduction

Animal sounds contain extensive information about community assemblage and ecosystem function across time and space (Burivalova et al. 2019, Sethi et al. 2020, Müller et al. 2023, Rasmussen et al. 2024), and this is particularly true for high-biodiversity environments such as tropical coral reefs, where most inhabitants produce and respond to sounds (Lobel et al. 2010, Tricas and Boyle 2014, Parmentier et al. 2021, Rice et al. 2022). In fact, the majority of fish species likely make sounds (Lobel et al. 2010, Tricas and Boyle 2014, Parmentier et al. 2021, Rice et al. 2022). In reef environments, high fish biodiversity and the presence of taxa within core functional groups play critical roles in reef health, resilience, and recovery (Duffy et al. 2016, Alsterberg et al. 2017, Hughes et al. 2018, Lefcheck et al. 2019, Lefcheck et al. 2021), and many reef fish species respond to environmental sounds (Simpson et al. 2016, Gordon et al. 2019). Previous studies suggest that the diversity, abundance, and behavior of these species could be detected by listening for species-specific sounds (Tricas and Boyle 2014), which could serve as key indicators of important functional groups, community integrity, or recovery processes (Kaplan et al. 2015, Tricas and Boyle 2021, Lamont et al. 2022b). One barrier has stood in the way – the technical difficulty or inability to decode the coral reefs’ complex soundscapes (Jarriel et al. 2024).

Globally, coral reefs and reef-based fisheries are in decline (MacNeil et al. 2015, Mouillot et al. 2016, Eddy et al. 2021). Improving long-term passive acoustic monitoring (PAM) capabilities for coral reefs can enable PAM to become an important complement to other technologies supporting conservation decision making (Apprill et al. 2023). For example, PAM provides finer spatial and temporal resolution than other approaches (e.g., eDNA, telemetry) (Mooney et al. 2020). PAM data can deliver valuable insights within ongoing monitoring efforts and inform actionable measures for adaptive management (Mooney et al. 2020, Lamont et al. 2022a, Apprill et al. 2023). PAM studies can target important protected or imperiled soniferous species (Schärer et al. 2012, Pyć et al. 2021), document ecosystem regime shifts, and examine soundscape composition patterns at the community level (Bertucci et al. 2016, Lin et al. 2021, Puebla-Aparicio et al. 2024). In fact, PAM may be better suited than other technologies to gather measurable, verifiable data on Key Performance Indicators (KPIs) relevant to innovative financing mechanisms for ocean conservation (Sumaila et al. 2023, International Capital Market Association 2024).

The fundamental challenge in realizing the utility of PAM for *in situ* fish assessments is that sounds from relatively few fish species have been identified (Looby et al. 2022, Rice et al. 2022), and the vast majority of recorded underwater biological sounds through PAM surveys cannot be attributed to a particular taxonomic group beyond “biological” or “fish” (Knudsen et al. 1948, Jarriel et al. 2024). Without taxonomic specificity (ideally to species level), PAM analysis can only yield measures of coarse spectral and temporal patterns. These measures have been used to derive indices that are *proxy* measures for biodiversity (Harris et al. 2016, Bradfer-Lawrence et al. 2023). While acoustic proxies have shown some correlation with on-the-ground diversity in some terrestrial ecosystems, they are not yet proving to reflect actual patterns of diversity in marine systems (Kaplan et al. 2015, Staaterman et al. 2017, Dimoff et al. 2021). Even when acoustic indices reflect biodiversity patterns, they remain computational abstractions of complex soundscapes and do not provide sufficient detail, such as direct species-level information, to drive conservation action through regulatory frameworks; conservation decision-making requires taxon-specific data (Godfray et al. 2004).

When species-specific sounds are known and available for comparison, we can directly measure species presence/absence, call type, and detection rates. Using presence/absence, we can decode the soundscape and implement species-informed biodiversity metrics, map species distributions, and detect indicator species (Fournet et al. 2019) and invasive species. When we know the sound types, we can map crucial behaviors such as spawning, territoriality, or feeding in space and time (Oestreich et al. 2024). To infer density and abundance requires knowledge of species identity and other information, such as detection probability (Marques et al. 2013). Therefore, species IDs enable the development of actionable metrics on which evidence-based adaptive management and policy decisions can be made (Fig. 1).

**Fig. 1.**
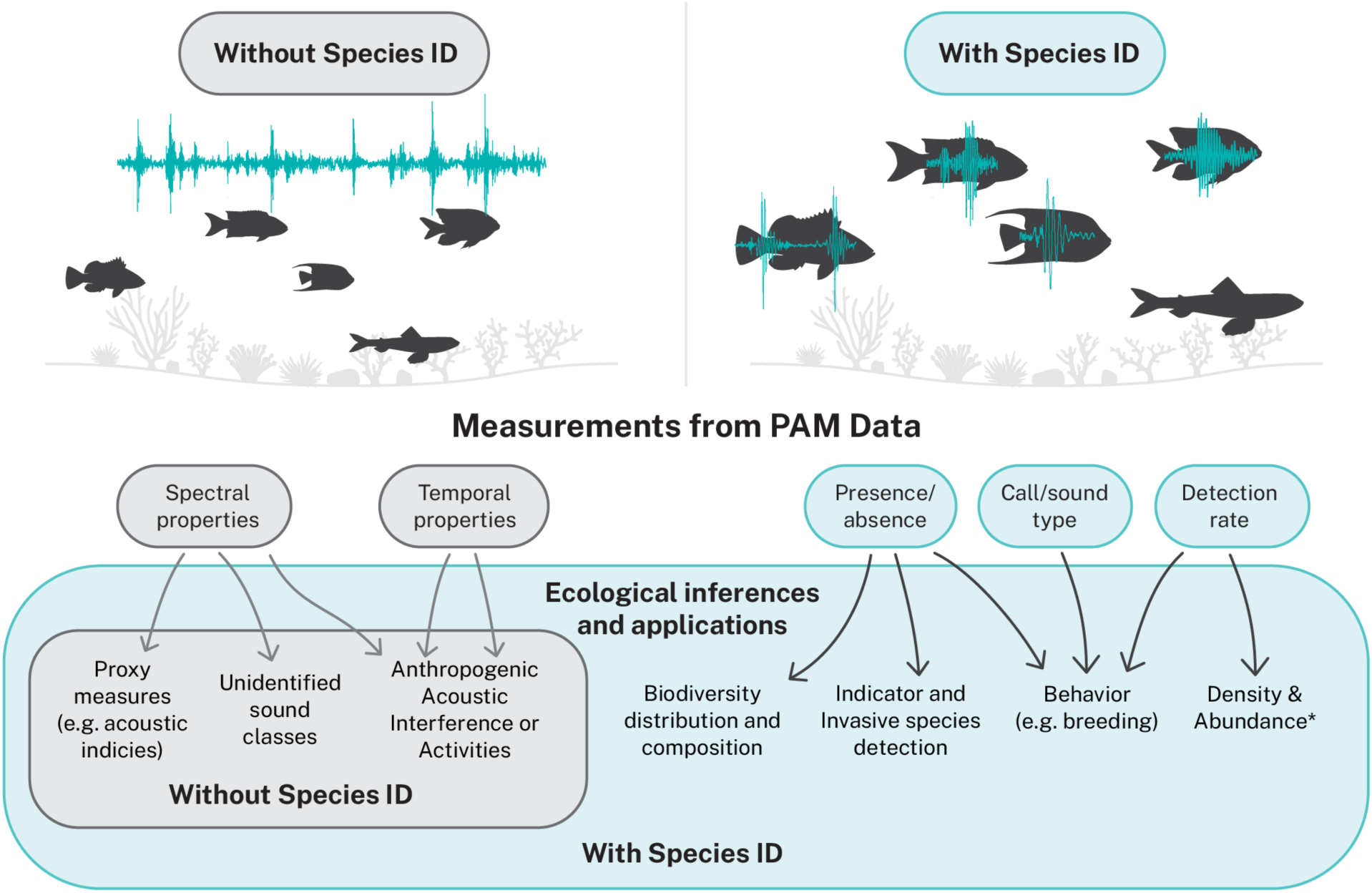
Conservation inferences from bioacoustics, both without and with species identities from sound. Without known species IDs, we can only measure the spectral and temporal properties of ambient sound. These can be processed into acoustic indices, examined for noise analysis, or clustered into sound types, limiting PAM studies to generalized categories such as biophony and anthrophony. Given species-specific sounds, we can directly create meaningful measures of biodiversity, detect indicator and invasive species, and monitor for behaviors. *Estimations of density and abundance require additional information, such as detection range and probability(Marques et al. 2013).

A lack of basic knowledge of species-specific fish sounds has severely limited opportunities for PAM in scalable and effective conservation approaches on coral reefs. For other taxonomic groups where species IDs are known– such as those of most species of birds, bats, cetaceans, and frogs– a long history of focal recordings has enabled the collection of a sufficient number of validated sounds (Kellogg 1960) to be used for the development of feature-based detection algorithms and ML methods (Kahl et al. 2021, Tuia et al. 2022). In systems where IDs are known (e.g., terrestrial, cetacean), these techniques are rapidly developing to enable long-term PAM monitoring systems supporting biodiversity conservation to function at large spatial and temporal scales (Gibb et al. 2019, Tuia et al. 2022, Rasmussen et al. 2024).

Evolutionary analysis suggests that 85% of fish families may make sounds, many of which could be used for acoustic monitoring (Rice et al. 2022). For example, in the region of focus for this study, the Caribbean Sea, of the 1122 reef-associated fish species(Robertson and Van Tassell 2023), 736 (66%) are in likely sonic families (Fig. 2; Supplement). A search of online sources (detailed in the Supplement) found records of 174 of these species with identified sounds, of which 132 have publicly available recordings. Only 33 species have recordings of free behaviors, 11 in nature. This means that less than 1.5% of Caribbean reef-associated fish species that likely make sound have available recordings from nature ascribed to them. Thus, recordings from a hydrophone on a Caribbean coral reef will have hundreds to thousands of likely fish sounds every minute, with variable call types suggesting a wide variety of sources. However, ascribing those sounds to individual species is exceedingly difficult (Kumpf 1964, Lobel 2005, Mouy et al. 2018, Mouy et al. 2023) because the species-specific vocalizations have not been identified. We have developed an approach to fill this knowledge gap and begun doing so for Caribbean fishes.

**Fig. 2.**
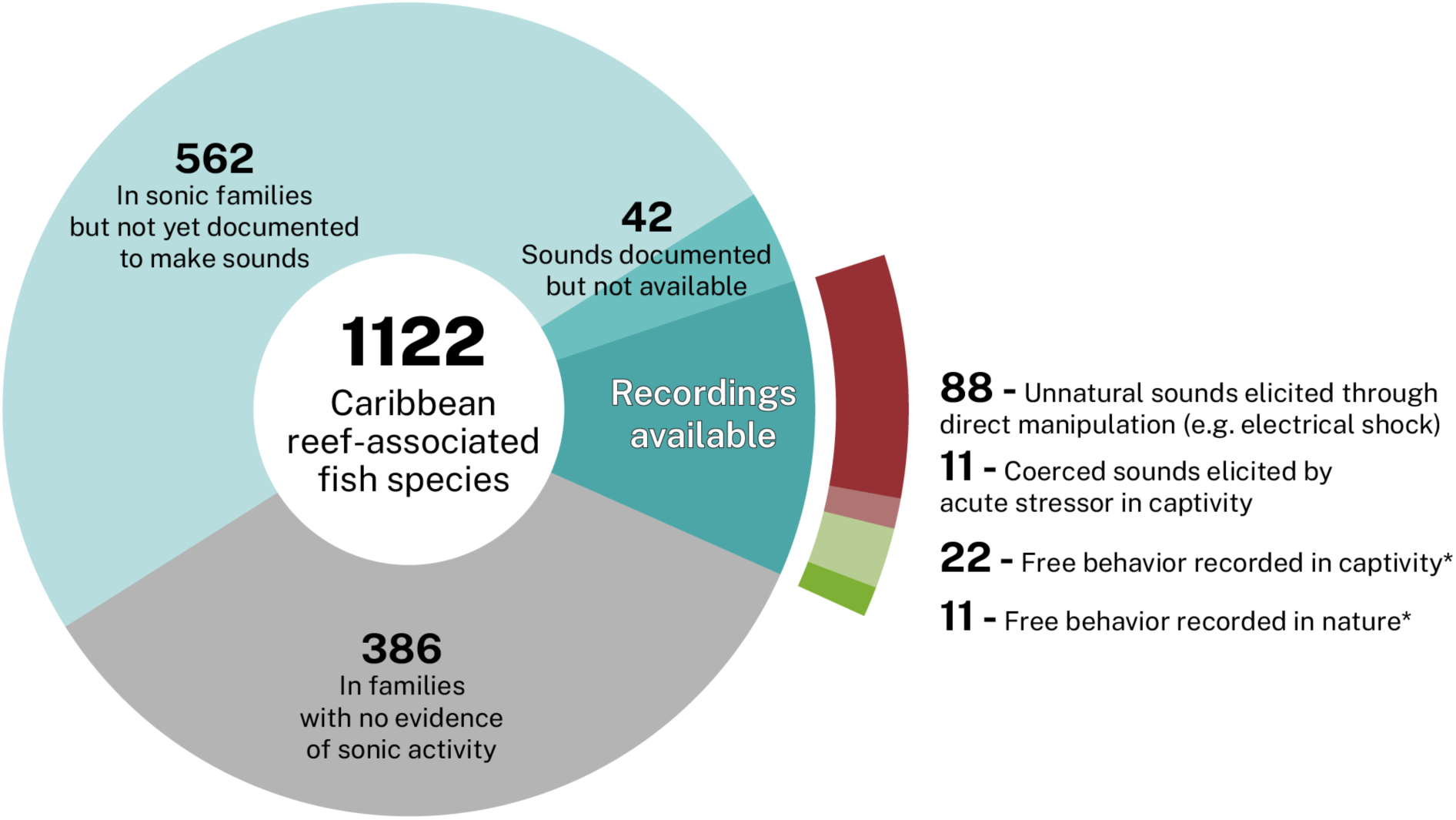
Summary of what is known about sounds for Caribbean reef-associated fishes. Of the total number of reef-associated fish in the Caribbean, 735 are in families likely to make sound [based on (Rice et al. 2022)]. A fraction of these species (174) have published evidence of acoustic behavior. Of these, we were able to find publicly available sound recordings for 132 species. We classified these recordings to grade their potential utility for PAM analysis. Most (88) of these sounds were elicited through direct manipulation of fish using techniques such as electric and manual stimulation. From top to bottom, the color of the “recordings available” relates to the utility of those recordings in analyzing in situ PAM recordings. We found only 11 Caribbean fish species with free behaviors recorded in nature. Details are in the supplementary text. *Species names and recording links are provided in Supplementary Table 1.

### Approaches to species identification

Despite fish sounds being common on coral reefs, ascribing those sounds to specific species is difficult. Underwater, humans can hear some fishes, but the ability of the human auditory system to localize sound underwater is minimal (Brandt and Hollien 1967, Casper and Babina 2022). Additionally, most fish sounds are made by cryptic stridulation or internal mechanisms without conspicuous visual correlates (Fine and Parmentier 2015). Because of this difficulty in making unaided observations, other methods have been used to try to ascribe sounds to fish species.

The most direct method to collect species-specific sounds is to record fish in a tank or other enclosure (Fig. 3a) (Fish and Mowbray 1970). While some fish are amenable to vocalizing in captivity (Rice and Bass 2009), for most species, natural behavior is uncommon under these circumstances, so many captive sounds have been recorded by subjecting the fish to unnatural stressors, including manual stimulation and electric shock (Fish and Mowbray 1970). These sounds might mirror natural distress calls but not the territoriality, courtship, or nest defense sounds that would be helpful in monitoring populations (Clark and Dunn 2022). In some cases, sounds have been recorded from habituated captive fish that may mirror a limited set of free behaviors (Rountree et al. 2006). However, the acoustic properties of tanks can make applying these recordings to PAM analysis challenging (Akamatsu et al. 2002). In nature, during group spawning aggregations or in rare cases where species are isolated, hydrophone recordings alone have enabled species identifications. Most of the 11 Caribbean field identifications were collected using this method, with only audio recordings (Locascio and Burton 2016). However, assigning identifications to sounds recorded in nature is fraught with the potential for misattribution (Sprague and Luczkovich 2001).

**Fig. 3.**
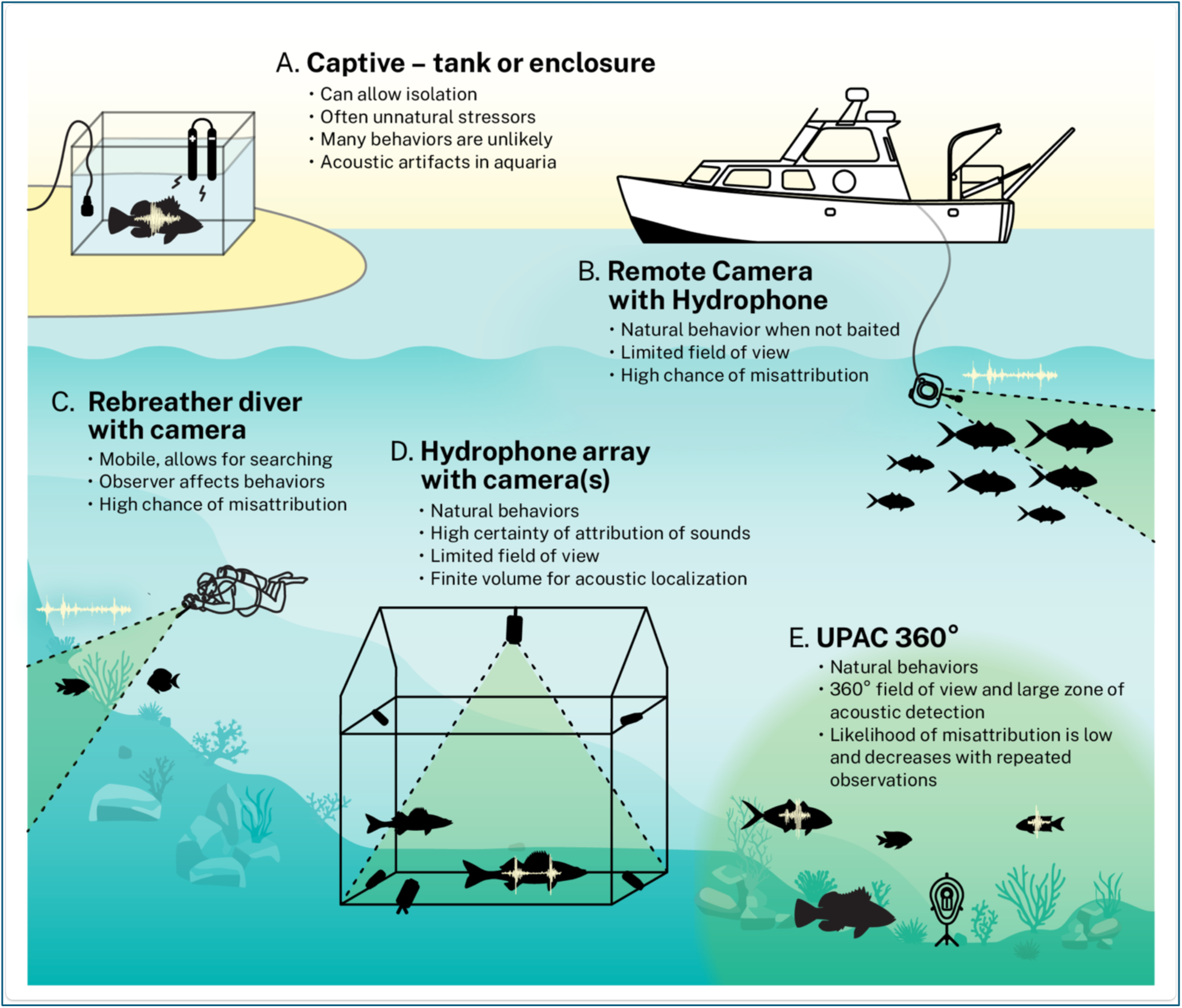
Methods of ascribing fish sounds to species using audio and video. It is challenging to ascribe sounds to fish species. Most studies use one of these methods. (**A**) Captive fish can be recorded in isolation, so attribution is clear, but natural behaviors involving sound production are rare, particularly for large species. (**B**) Remote cameras with hydrophones can capture natural behavior but with a narrow FOV and a high probability of misattribution. (**C**) Rebreather divers have a flexible FOV but retain similar limitations. (**D**) A method to increase the certainty of attribution uses hydrophone arrays to ensure that a sound is coming from the location of a visible fish. (**E**) our approach uses a 360° unbounded FOV with a compact hydrophone array, providing a small footprint, low impact on fish behavior, and a large zone of detection.

In a diverse environment, adding synchronous video recordings can enable verification of identifications and provide greater certainty of species isolation. The coupling of video and audio has been accomplished with remote cameras (Mann et al. 2010, Banse et al. 2024) and by divers (Lobel 1992). Scuba divers have the flexibility to search for soniferous behaviors, and the most successful diver recordings have used closed-circuit rebreathers to minimize diver disturbance (Lobel 2005, Tricas and Boyle 2014). However, underwater diver time is limited, so recording with divers is more time-limited than fixed autonomous installations. There is a high chance of misattribution for both remote cameras and divers because they use directional cameras and omnidirectional hydrophones. Therefore, the individual fish within the limited FOV may not be the source of the sounds recorded (Fig. 3b, c) (Mouy et al. 2018, Mouy et al. 2023).

To address the potential for misattribution, one method has paired remote cameras with a hydrophone array for acoustic localization (Fig. 3d) (Mouy et al. 2023). The multichannel recordings are used for acoustic localization. Sounds are only ascribed to individuals when the localization points to a fish within the FOV. Sometimes, localization is precise enough to distinguish between sources when more than one fish is visible within the FOV. However, the cameras used in this method have a limited directional FOV, and localizations that are sufficiently precise to ascribe sounds can only be made in small regions (Mouy et al. 2023).

Here, we describe and apply a novel technique (Fig. 3e) to a remote camera solution that avoids the pitfalls of previous methods, such as limited FOV, human disturbance, and misattribution. This technique is far more productive than previous methods, allowing the identification of many sounds each day of recording. Thus, we demonstrate that a scalable and generalizable fish sound library can be built. Such sounds and recordings can improve PAM analysis and deliver critical insight into community assemblage and ecosystem function.

## Materials and Methods

### Hardware Details

Our acoustic array (Fig. 4a) is modeled on an earlier version of a tetrahedral underwater ambisonics array, which successfully tracked a scuba diver in a pool (Delikaris-Manias et al. 2018). Due to the uniform placement of the hydrophones, the performance of the algorithms is independent of the direction of the sound source. We paired the geometry of that array with an extruded acrylic frame that holds the four hydrophones (a prototype of the A5 hydrophone from Aquarian Audio and Scientific, Anacortes, WA) in a tetrahedral pattern. The frame is designed to position the array around a 360° camera (Insta 360, X3, Shenzhen, Guangdong, China) with an underwater housing. The frame is not visible because objects close to the camera in the plane between the lenses are not visible in the composite image. The one hydrophone and cable in the FOV are visible, but because of the clear acrylic, the small size, and the orientation of the hydrophone, its visual footprint is minimized (Fig. 4b).

**Fig. 4.**
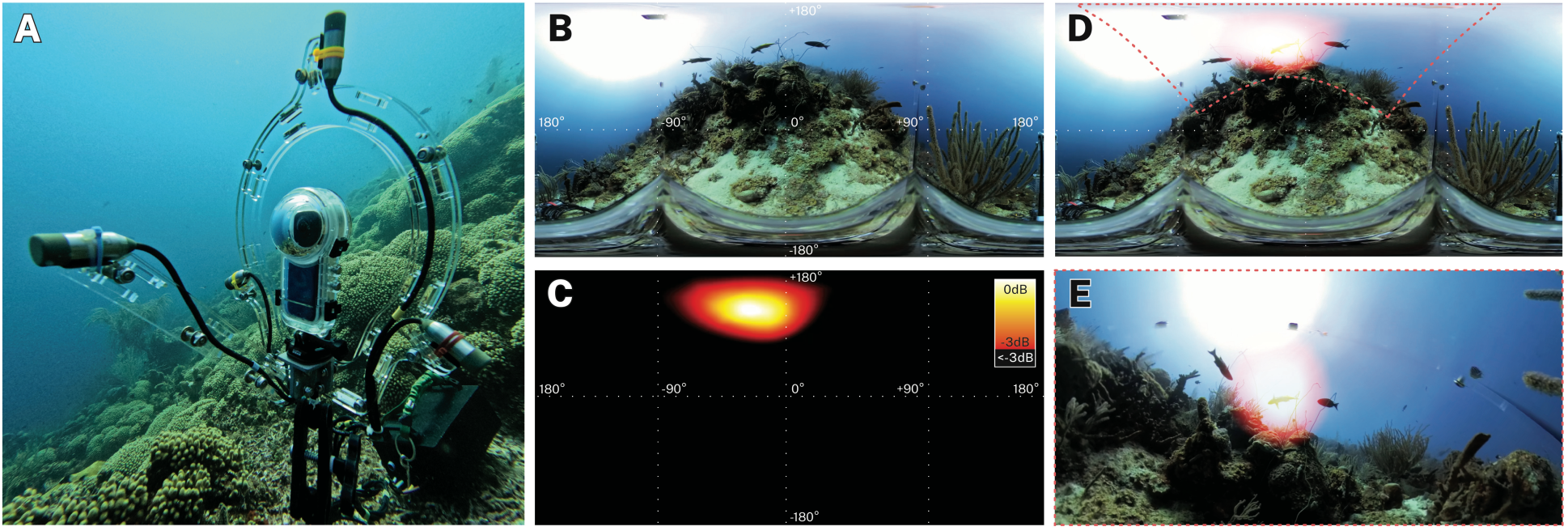
Conceptual overview of an underwater passive acoustic camera, combining spatial audio with 360° video and data visualization. (**A**) The UPAC-360 comprises a sensor package of four hydrophones in a compact tetrahedral array with a commercial 360° camera at the array’s center. (**B**) The equirectangular (EQR) image from the camera after processing (stitching) the images from its two sensors. The obstruction at the bottom of the frame is the camera housing. (**C**) The powermap visualization of the sound field energy distribution at the moment of a fish sound. We display the top 3dB of this map on the same EQR projection. (**D**) We overlay the powermap visualization onto the video. (**E**) We reframe the image to “look” toward the detection of an overlapping sound source.

The videos were recorded at 5760 x 2880 resolution at 30 frames per second in automatic exposure mode. The audio was recorded on a Zoom F6 field recorder (Zoom Corporation, Tokyo, Japan) set to 48 kHz sample rate and 32-bit depth, with the channel gains locked to record in Ambisonic A format. The audio recorder was housed in a pressure housing (Blue Robotics Inc., Torrance, CA).

### Field Effort

The recording system was deployed at a diversity of coral reef sites at depths of 3-45 m off the Curaçao Sea Aquarium in Bapor Kibrá, Willemstad, Curaçao (12.084°N, 68.896°W) in June-July 2023, December 2023, and July 2024. Here, we are reporting on the analysis of 20 hours of recordings from June 2023. Analysis of the remaining materials is ongoing. The June recordings were made over four days in six locations.

### Analysis Methodology

Raw video files were exported as 5760 x 2880 EQR files using Insta360 Studio (Fig 4B). We then aligned audio and video in DaVinci Resolve v. 19 (Black Magic Design, Freemont, CA, USA), using diver-generated and natural sounds as reference points. We excised the audio sections that align with each camera deployment for further analysis. We converted the four-channel audio captured with the hydrophone array to first-order ambisonic signals through a spherical harmonic transformation adopted for the hydrophone array configuration(McCormack et al. 2018).

The fish sound detection was performed manually familiar with fish sounds inspecting spectrograms. We then utilized the ambisonic signals to visualize the energy of the soundfield in Adobe Audition (Adobe, San Jose, CA, USA) using the SPARTA VST toolbox to inspect if there was an energy peak at the same time (McCormack et al. 2018). We marked these calls for subsequent analysis that uses a combination of Multiple Signal Classification (MUSIC)(Schmidt 1986) and Cross-Pattern Coherence (CroPaC) algorithms (Delikaris-Manias and Pulkki 2013) to visualize the soundfield energy. The sound analysis was performed in the frequency range of 200-2500 Hz, encompassing most known fish sounds. We normalized each section to its energy peak, and the soundfield visualization depicts the highest 3 dB (Fig 4c). We screen-composited these maps over the camera’s video in DaVinci Resolve and adjusted opacity to allow the video to show through (Fig. 4d). To focus on the event, we reframed 360° video into standard framing using the Karta VR plugin V.5.7 (Kartaverse, https://kartaverse.github.io/) (Fig. 4d).

### Assigning sounds

Our recordings have hundreds of likely fish sounds every hour, but we cannot assign most of them to individuals or species. The reasons are: 1) Many sounds recorded by the array come from sources beyond our visual range. The visual range is limited by light level, water visibility, and camera resolution. 2) The topography and coral coverage obscure many fish wholly or partially. 3) Many fish are visually cryptic during the day. 4) Color variations mask distinguishing features between closely related species. 5) The spatial detection area for these sounds overlaps with more than one species. However, we can overcome some of these challenges using additional evidence. For example, sound production often coincides with conspicuous or subtle movements, allowing us to discriminate between individuals. While single events of sound production may be difficult to ascribe to species with certainty, multiple examples lend weight to evidence.

### Integrating UPAC-360 Data with PAM Recordings

To demonstrate how validated sounds through the UPAC-360 approach could be used in traditional PAM approaches, we deployed the UPAC-360 next to a bottom-mounted archival acoustic recorder (SoundTrap ST600, Ocean Instruments, Auckland, NZ) sampling at 48 kHz. The SoundTrap was mounted to an old car tire found on the reef. A random five-minute segment of the audio from the SoundTrap was chosen for annotation. Identifiable sounds to the lowest taxonomic resolution(Robertson and Van Tassell 2023) were annotated through visual spectrographic analysis in Raven Pro 1.6.5. Unidentifiable sounds were manually sorted into classes based on the similarity of distinct spectral and temporal features.

## Results

We created a novel approach to ascribing sounds to fish species, which we describe as an omnidirectional (360°) Underwater Passive Acoustic Camera (UPAC 360°). It combines a four-element hydrophone array yielding underwater ambisonics recordings (Fig. 4a) with concentric 360° video (Fig. 4b), coupling commercially available technologies with publicly available spatial audio techniques. Visualizations of the detected sound field (Fig. 4c) are overlaid on the video (Fig. 4d), revealing the source of the received acoustic signal (Fig 4e). In this example, all three fish within the zone of detection are *Clepticus parrae*, so we ascribe the sound to that species. Using behavioral cues from the video, we can further ascribe the sound to the fish at the center.

The UPAC-360, with spatial audio algorithms, can localize many sounds, even when the reef is crowded with activity (Figs. 5, S1). Within this video frame are more than 121 individuals from at least six species. The detection zone overlaps only one individual, which can be identified as a *Stegastes partitus* (Bicolor Damselfish) just before and after the rapid dive this species does while making this sound.

**Fig. 5.**
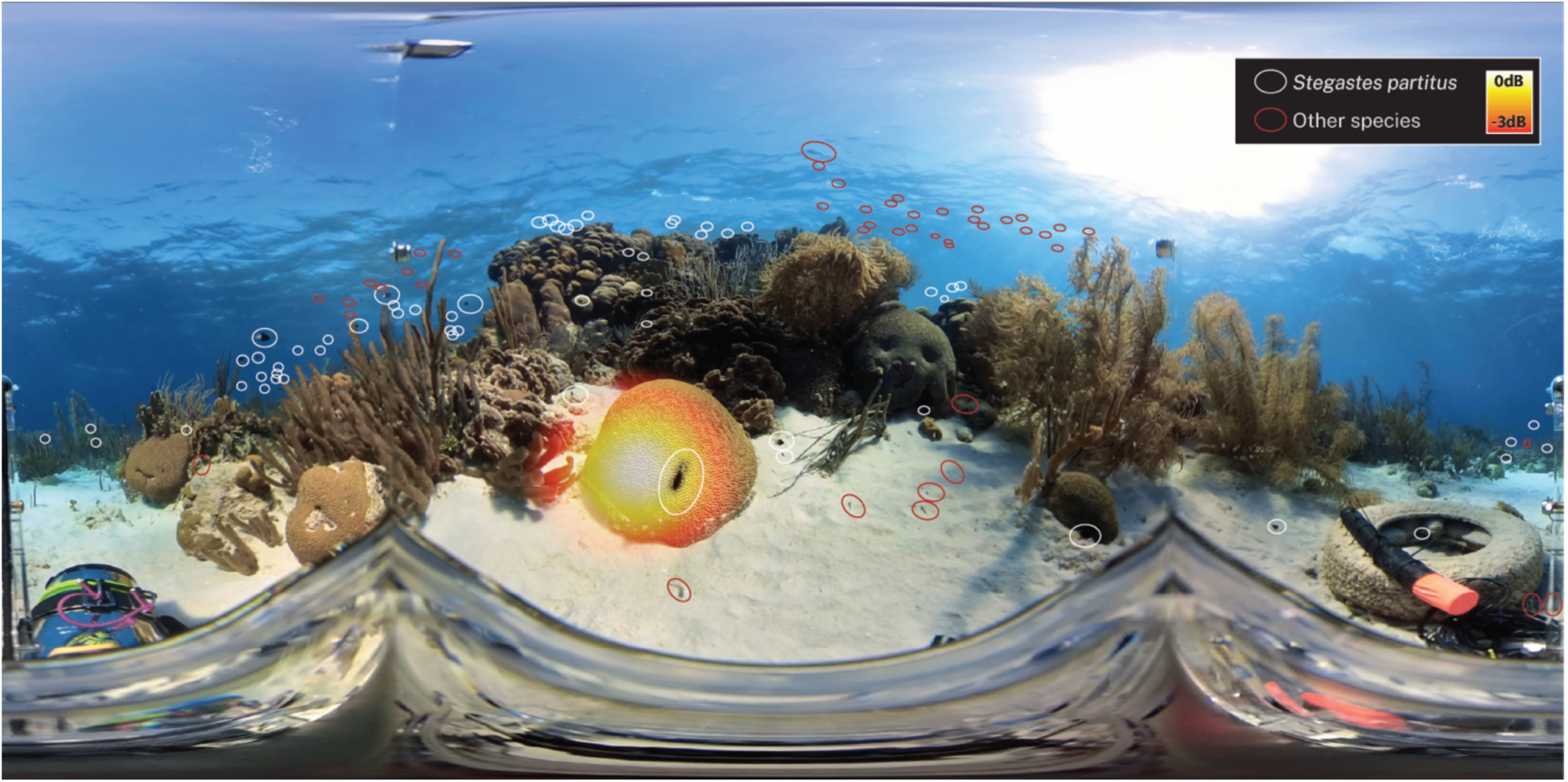
Ascribing sounds to individuals. The UPAC-360 can isolate which individual is the sound source in a coral reef. Here, we identified at least 121 individuals from six species, including 58 *Stegastes partitus,* greater than 53 *Azurina multilineata*, 1 *Lactophrys bicaudalis*, 1 *Halichoeres garnoti*, 2 *Canthigaster rostrata*, and 5 from the family Gobiidae. The 3 dB zone overlaps only one individual *S. partitus.* The visual ID is based on video before and after the soniferous event (**Supplementary Movie 1**). The bottom left shows the Blue Robotics housing for the recorder. In the bottom right corner is the PAM recorder which is the source of the data for Fig 7.

Using this technique, we have ascribed sounds to species. Figure 6 (and Supplementary Movie 2) shows examples of sounds ascribed to individuals from six species (*Heteropriacanthus cruentatus*, *Clepticus parrae*, *Ocyurus chrysurus*, *Scarus taeniopterus*, *Azurina multilineata*, and *Stegastes partitus*). We documented both active (intentional) and passive (non-intentional) sounds. The sounds from *H. cruentatus*, *C. parrae*, and *O. chrysurus* were active sounds from isolated individuals. The *S. taeniopterus* sound is an epiphenomenon of grazing activities, an example of a passive (non-intentional) sound. In the *Azurina multilineata* and *Stegastes partitus* examples, there is more than one individual in the detection zone, but all are the same species.

**Fig. 6.**
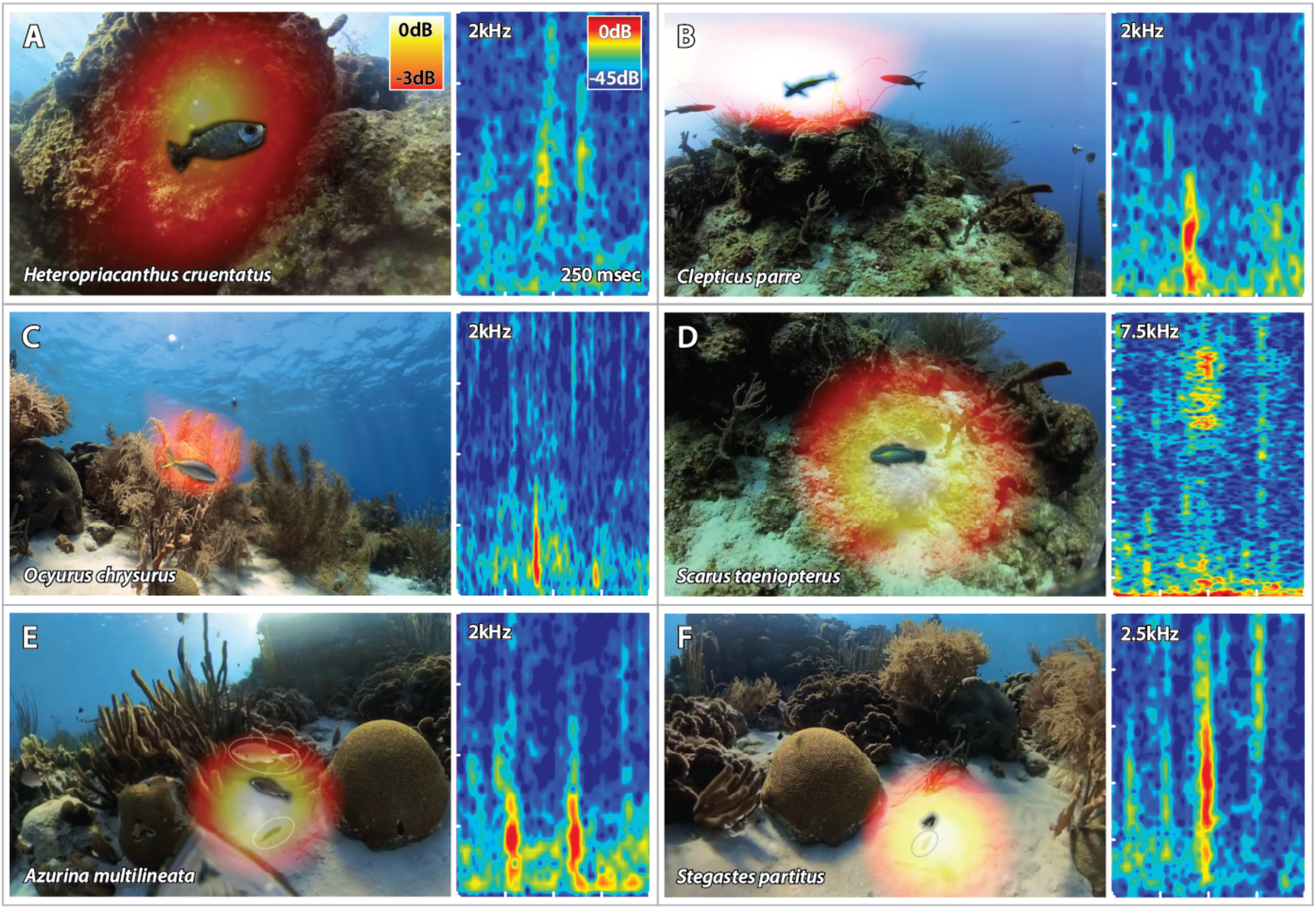
Ascribing sounds to species. The sounds of (**A**) *Heteropriacanthus cruentatus*, Glasseye Snapper, (**B**) *Clepticus parrae*, Creole Wrasse, (**C**) *Ocyurus chrysurus*, Yellowtail Snapper, (**D**) *Scarus taeniopterus*, Princess Parrotfish, (**E**) *Azurina multilineata*, Brown Chromis, and (**F**) *Stegastes partitus*, Bicolor Damselfish. For (E) *A. multilineata* and (F) *S. partitus,* there is more than one individual in the zone of detection (white ovals), but all are the same species. We can distinguish which individual made the sound by behavior (Supplementary Movie 2). The powermap data layer is masked to make the individual visible in this illustration. Spectrograms to the right of each image are made on the 0-order spherical harmonic of the ambisonic audio using Raven Pro 1.6.5, Hanning window, 1024-point FFT, 75% overlap.

The combination of 360° video with underwater spatial audio in an integrated visualization allows for a scalable ability to ascribe sounds to fish species in a behaviorally relevant context. From UPAC-360 deployments over four days on reefs in Curaçao, we have identified the sounds of 40 species of Caribbean reef-associated fishes (Table 1) belonging to 17 families. None of these have publicly available natural sounds. 18 of these fish species have been shown to be soniferous under duress in captivity (†), and four have been documented but lack publicly available recordings (‡, see Fishsounds.net). The remaining species (19) have never been recorded in any context. The sounds we have collected that have a high degree of certainty will be publicly available through our website (library.fisheyecollaborative.org, to be opened at the time of final publication) for future qualitative or quantitative comparison to previous and future acoustic collections from Caribbean soundscape studies. With the ongoing application of the technique, we expect the number of acoustic examples to include new species and specimens to be regularly added to the open-access library.

**Table 1.** List of species with ascribed sounds. Media specimens for 40 species will be available in the FishEye Collaborative Library (library.fisheyecollaborative.org) at the time of final publication. They will be found on their species’ pages and by their catalog numbers. Up to five high-confidence representative recordings will be included for each species.

| Family | Scientific Name | Common Name | Catalog Numbers |
| --- | --- | --- | --- |
| Acanthuridae | *Acanthurus coeruleus* † | Blue Tang | FEC-00006<br>FEC-00007<br>FEC-00008<br>FEC-00009<br>FEC-00010 |
|  | *Acanthurus tractus* | Northern Ocean Surgeonfish | FEC-00011<br>FEC-00012 |
| Aulostomidae | *Aulostomus maculatus* | Atlantic Trumpetfish | FEC-00013<br>FEC-00014<br>FEC-00015 |
| Carangidae | *Caranx ruber* † | Bar Jack | FEC-00026<br>FEC-00027<br>FEC-00028<br>FEC-00029<br>FEC-00030 |
| Chaetodontidae | *Chaetodon capistratus* | Foureye Butterflyfish | FEC-00109<br>FEC-00110<br>FEC-00111<br>FEC-00112 |
| Haemulidae | *Anisotremus virginicus* † | Porkfish | FEC-00103<br>FEC-00104<br>FEC-00105<br>FEC-00106<br>FEC-00107 |
|  | *Brachygenys chrysargyrea* | Smallmouth Grunt | FEC-00024 |
|  | *Haemulon flavolineatum* † | French Grunt | FEC-00038 |
|  | *Haemulon vittatum* | Boga | FEC-00113<br>FEC-00114 |
| Holocentridae | *Myripristis jacobus* † | Blackbar Soldierfish | FEC-00059<br>FEC-00060<br>FEC-00061<br>FEC-00062<br>FEC-00063 |
| Kyphosidae | *Kyphosus bigibbus* | Darkfin Sea Chub | FEC-00050<br>FEC-00051 |
| Labridae | *Bodianus rufus* | Spanish Hogfish | FEC-00108 |
|  | *Clepticus parrae* | Creole Wrasse | FEC-00032<br>FEC-00033<br>FEC-00034<br>FEC-00035<br>FEC-00036 |
|  | *Halichoeres garnoti* | Yellowhead Wrasse | FEC-00039<br>FEC-00040<br>FEC-00041<br>FEC-00042<br>FEC-00043 |
|  | *Thalassoma bifasciatum* | Bluehead Wrasse | FEC-00102 |
| Lutjanidae | *Lutjanus apodus* † | Schoolmaster | FEC-00115<br>FEC-00116<br>FEC-00117<br>FEC-00118 |
|  | *Lutjanus griseus* † | Gray Snapper | FEC-00053 |
|  | *Lutjanus mahogoni* ‡ | Mahogany Snapper | FEC-00119<br>FEC-00120<br>FEC-00121<br>FEC-00122 |
|  | *Ocyurus chrysurus* † | Yellowtail Snapper | FEC-00064 |
| Mullidae | *Mulloidichthys martinicus* † | Yellow Goatfish | FEC-00058 |
| Ostraciidae | *Lactophrys triqueter* † | Smooth Trunkfish | FEC-00052 |
| Pomacentridae | *Abudefduf saxatilis* † | Sergeant Major | FEC-00001<br>FEC-00002<br>FEC-00003<br>FEC-00004<br>FEC-00005 |
|  | *Microspathodon chrysurus* | Yellowtail Damselfish | FEC-00054<br>FEC-00055<br>FEC-00056<br>FEC-00057 |
|  | *Stegastes diencaeus* ‡ | Longfin Damselfish | FEC-00087<br>FEC-00088<br>FEC-00089<br>FEC-00090<br>FEC-00091 |
|  | *Stegastes partitus* ‡ | Bicolor Damselfish | FEC-00092<br>FEC-00093<br>FEC-00094<br>FEC-00095<br>FEC-00096 |
|  | *Stegastes planifrons* ‡ | Threespot Damselfish | FEC-00097<br>FEC-00098<br>FEC-00099<br>FEC-00100 |
|  |  |  | FEC-00101 |
|  | *Azurina cyanea* | Blue Chromis | FEC-00016<br>FEC-00017<br>FEC-00018 |
|  | *Azurina multilineata* | Brown Chromis | FEC-00019<br>FEC-00020<br>FEC-00021<br>FEC-00022<br>FEC-00023 |
| Priacanthidae | *Heteropriacanthus cruentatus* † | Glasseye Snapper | FEC-00044<br>FEC-00045<br>FEC-00046<br>FEC-00047<br>FEC-00048 |
| Scaridae | *Scarus iseri* † | Striped Parrotfish | FEC-00065<br>FEC-00066<br>FEC-00067<br>FEC-00068<br>FEC-00069 |
|  | *Scarus taeniopterus* | Princess Parrotfish | FEC-00070<br>FEC-00071<br>FEC-00072<br>FEC-00073 |
|  | *Scarus vetula* † | Queen Parrotfish | FEC-00074<br>FEC-00075 |
|  | *Sparisoma aurofrenatum* † | Redband Parrotfish | FEC-00076<br>FEC-00077<br>FEC-00078<br>FEC-00079<br>FEC-00080 |
|  | *Sparisoma rubripinne* † | Yellowtail Parrotfish | FEC-00081 |
|  | *Sparisoma viride* † | Stoplight Parrotfish | FEC-00082<br>FEC-00083<br>FEC-00084<br>FEC-00085 |
| Sciaenidae | *Eques punctatus* | Spotted Drum | FEC-00037 |
| Serranidae | *Cephalopholis fulva* † | Coney | FEC-00031 |
|  | *Hypoplectrus chlorurus* | Yellowtail Hamlet | FEC-00049 |
| Tetraodontidae | *Sphoeroides spengleri* | Bandtail Puffer | FEC-00086 |
|  | *Canthigaster rostrata* | Sharpnose Puffer | FEC-00025 |

To demonstrate how acoustic identifications can be used to decipher soundscape recordings, we analyzed a representative five-minute section of a PAM recording collected on Dec 17, 2023, on the co-located SoundTrap (seen in Fig. 5). We identified and manually annotated 675 fish sounds in this five-minute section. Using our library of reference sounds, we were able to determine distinct characteristics such as frequency range, number of pulses, and pulse rate, along with a qualitative comparison with the acoustic library for some of the most common species on the reef. We used this information to identify unique species-specific sounds, such as multi-pulsed calls produced by *Clepticus parrae, Stegastes partitus*, and *Azurina multilineata,* with the latter typically occurring at higher frequencies and containing fewer pulses. Other Pomacentridae species produce similar pulsed sounds in similar frequency ranges; therefore, in cases where we were uncertain about any specific species, we identified these sounds to the family level. There were many sounds that we could not identify, largely low-frequency, extremely quiet, or harmonic sounds produced by fish that we either do not yet have a substantial set of sounds for or have not yet recorded. In these cases, we grouped the unknowns into 13 distinct acoustic classes based on similarity (Fig. 7). The wide range of unknowns shows the incredible diversity in sounds and likely cryptic species in even a short excerpt of a PAM recording. Given the long-term recording capabilities of PAM devices and our ability to continuously grow a robust library of confirmed fish sounds, it is likely that these unknown sounds will become identifiable, and the number of species that can be confirmed through PAM recordings will grow.

**Fig. 7.**
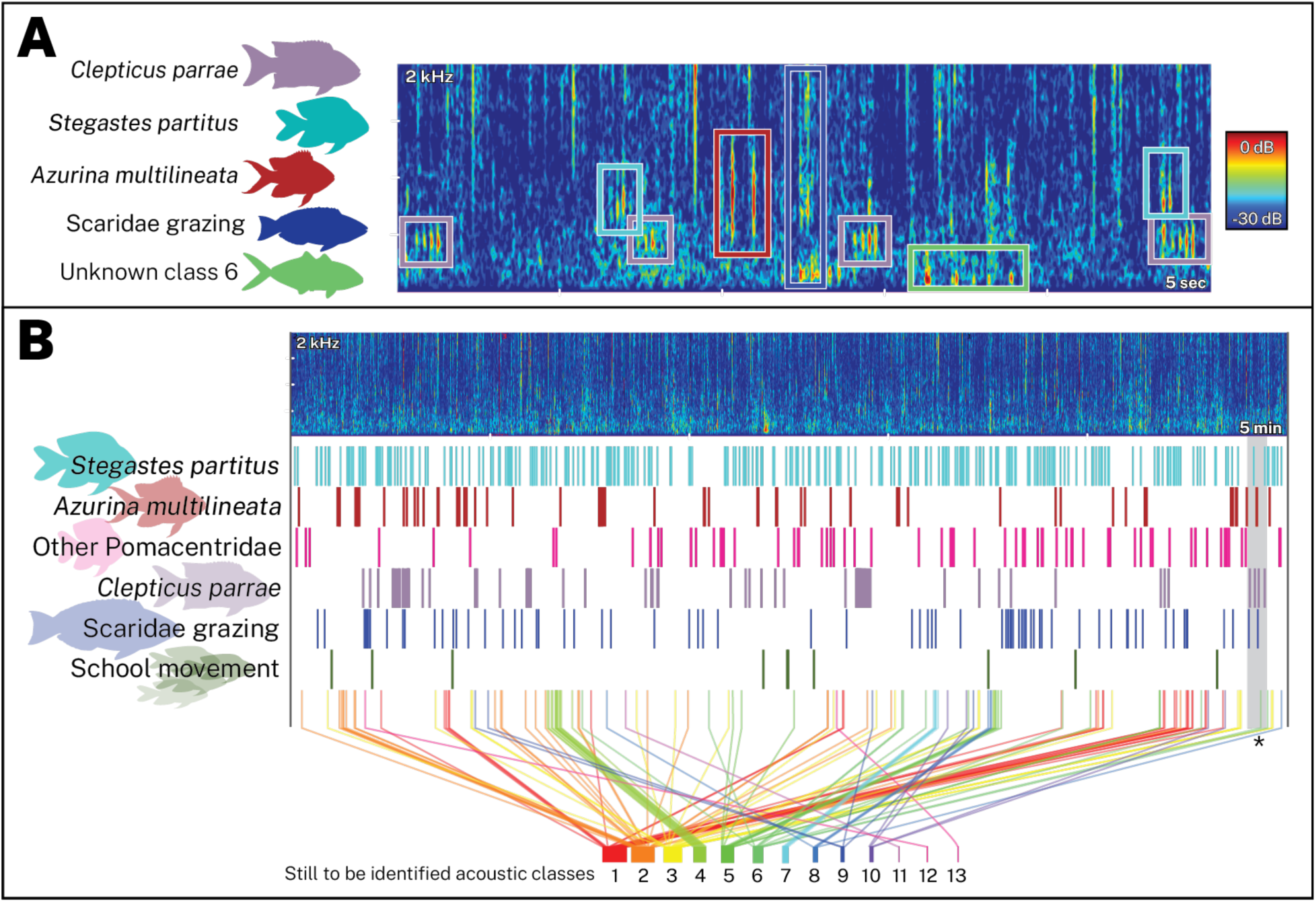
Identification of species-specific sounds in archival passive acoustic recording. (**A**) Spectrogram of a representative 5 s segment of sound, showing the labeling of five classes of sounds, three to species level (*C. parrae*, *S. partitus*, *A. multilineata*), parrotfish feeding sounds, and a class of sounds of an unidentified species. (**B**) Spectrogram of representative 5 min of archival audio with three classes of sound to species, two classes to family, one to a behavioral category, and 13 unidentified sound classes, with the occurrence in time labeled by a colored bar. *The grey box corresponds to the 5 s spectrogram in panel (A). The sound was recorded on a SoundTrap ST600 at 8 kHz, 16-bit depth. Spectrograms are made using Raven Pro 1.6.5, Hanning window, 256-point FFT, and 75% overlap.

## Discussion

We have developed an approach that enables matching species with sounds in crowded and complex systems. We have demonstrated the capabilities of this approach for Caribbean reef-associated fishes, but the technique applies to reefs around the world as well as in other environments with clear water and fish diversity. We have identified the sounds of many new species, but many more sonic species remain. Therefore, much more sampling and analysis need to be done.

### Using reference sounds for analysis

Manually cross-referencing sounds in a PAM recording with validated IDs (e.g., see Fig. 7) is labor-intensive and not scalable. Therefore, automated detector algorithms are required to extract patterns from months or years of PAM data. These detectors can either be traditional feature-based or ML-derived, with ML detectors offering the most promise for fine-scale distinguishing between the sounds many fish make, for which species classification is more challenging than for taxa like birds or whales. The development of both approaches has been stymied for fishes by the lack of sufficient ground-truthed acoustic data sufficient for species identification (Yassir et al. 2023). For ML, the lack of data annotation and class imbalance has led to a reliance on unsupervised approaches, which leads to low classification accuracy and generalizable applicability (Mouy et al. 2024). Traditional models have also struggled to advance without sufficient ground truth information or the capacity to support multiclass supervised approaches, which have proven effective for other taxa (van Merriënboer et al. 2024), particularly birds. Our acoustic library provides this ground truthing of species identification and behavioral context, addressing these shortcomings for both types of models, thereby assisting in the model development that will be necessary to gain the critical insights that can come from long temporal and high spatial resolution datasets (Parsons et al. 2022).

Only some sounds are common enough on reefs to avoid class imbalance. In these cases, researchers can use identified sounds from reference collections to qualitatively or quantitatively identify the same sounds in their own recordings or publicly available PAM survey data, allowing them to start aggregating validated sounds to build a training dataset for ML models. Publicly accessible reference libraries, such as the one we are building and other global efforts (Parsons et al. 2022), will play a critical role in broadening the accessibility of sounds and enable multiple groups to use publicly available data to analyze their sounds.

However, building a larger set of UPAC-360 validated sounds for each species will be necessary, as will a broader spatial and geographical application of this approach. Little is known about how fish sounds change with factors such as locality, temperature, and depth, and similar questions remain for reef ecosystems in the Indo-Pacific. As with other taxa, sounds may vary by population (dialects) (Parmentier et al. 2005). The sounds may point to cryptic species divergence (Parmentier et al. 2022), which would also compromise the geographic applicability of detectors. Therefore, the ID work must be applied to more locations and for more extended periods to amass larger validated sets. By building a robust and diverse sound repository, we can address data gaps, enabling the development of machine-learning tools that support more effective and scalable monitoring (Ditria et al. 2022).

### Behavioral context - more than just species ID

The primary purpose of building a publicly available library of reference fish sounds is to enable decoding soundscape recordings made with PAM devices (Parsons et al. 2022). Most conservation-relevant ecological inferences and applications (e.g., presence/absence for direct measures of biodiversity, indicator, and invasive species detection (Fig. 1)) can be measured from PAM using just acoustic IDs. However, our UPAC-360 method captures more than just sonic IDs; we also capture a record of the behavioral context. The system is semi-autonomous, running unattended for hours once deployed, allowing us to capture natural behaviors (e.g., agonistic, mating/territorial displays, grazing (Supplementary Movie 3). Because the device has a 360° FOV, we often see the entire context of acoustic events without losing track of individuals that come in and out of the FOV of traditional video. Thus, we gain insights into behaviors that are largely absent from previous studies, few of which have any confirmed behavioral context for their sounds (Looby et al. 2023a). Changes in fish behavior may serve as a leading indicator of ecosystem regime shifts or outcomes of conservation interventions.

### PAM with species ID can address data deficiencies

Marine conservation suffers from significant data deficiencies, making it challenging to formulate actionable policies to address biodiversity loss and coral reef decline (Steneck et al. 2018, Mouquet et al. 2024), resulting in mismanaged Marine Protected Areas (MPAs) (Grorud-Colvert et al. 2021). Coupling technology with traditional methods has proven to progress conservation action (Madin et al. 2019, Muenzel et al. 2024). International organizations, such as the United Nations, the International Union for Conservation of Nature (IUCN), the Convention on Biological Diversity (CBD), and the World Bank, have established ambitious conservation goals and metrics (Colglazier 2015). However, indices to meet these goals are under debate and often unattainable with current technologies, requiring novel approaches (Diaz-Sarachaga et al. 2018). Our methodology and application highlight the power and potential growth of conservation approaches by integrating acoustic monitoring with ongoing efforts, providing tools to evaluate ecosystem health through actionable metrics (Fig. 1).

Species-specific identifications and passive acoustic recording techniques offer a promising solution to further close the gap of enabling continuous, scalable, and adaptive monitoring (Grorud-Colvert et al. 2021). Conventional monitoring strategies often lack the spatial and temporal resolution necessary for actionable change. While eDNA monitoring is increasingly seen as the primary tool to address data deficiencies (Muenzel et al. 2024), PAM with verified ID has greater spatial and temporal precision, allowing the detection of specific events or places that need protection. This precision is also required to monitor specific impacts, such as spill-over effects of MPAs on surrounding fisheries, as well as the range of those effects. This type of precision and scale are going to be required as KPI if the types of large-scale financing efforts that are growing in prevalence are to have their intended scale of impact.

Rates of biodiversity and habitat loss are at an unprecedented high (Dirzo et al. 2014), and conventional monitoring approaches are insufficient to address these challenges (Langhammer et al. 2024). Classified sound data offer the resolution needed to answer questions about species distributions, abundances, habitat requirements, and threats. Long-term acoustic monitoring provides spatial, temporal, and historical insights that enable global applications. Moreover, acoustic sensors are becoming increasingly affordable and widely implemented across conservation sites. Advancements in technology, such as distributed acoustic systems, are further enhancing spatial resolution beyond single-sensor applications. These are factors that are leading to the growth of PAM in terrestrial and marine habitats around the world. Adding verified IDs to PAM techniques can similarly make acoustics a transformative tool for the conservation of the coral reefs and other nearshore fish habitats that support hundreds of millions of lives, contain a disproportionate level of biodiversity and suffer a disproportionate share of the impacts of climate change and direct human degradation.

## Acknowledgments

The authors wish to thank: Cooper Nichols designed the acrylic bracket. Robb Nichols of Aquarian Audio made the prototype hydrophones available for our use. Charlie Dantzker provided design and engineering support. Brayden Zee developed analysis software. In the field, we were given invaluable support by Adrien “Dutch” Schrier and the staff of the Curaçao Sea Aquarium and Substation Curaçao including Tafari Bakmeijer, Tico Cristiaan, Manuel Jove, Loreene Schenk, Jordy Stolk, and Joel Tjong-A-Tjoe. By arrangement with the Sea Aquarium, Aldemar Rodriguez served as our field technician whose experience on the local reef proved invaluable. We’d also like to thank Carole Baldwin and Matthew Girard from the National Museum of Natural History at the Smithsonian Institution for their involvement in our field operations. Heather Dantzker gave significant editorial feedback on the manuscript. Thanks to Aaron Adams, Brent Miller, and Will Palmer, our volunteer web team who wrote, designed, and developed the online library.

## Funding

Schmidt Marine Technology Partners, Schmidt Family Foundation (MSD, ANR, VP)

Oceankind, LLC (ANR, MSD)

NSF Predoctoral Fellowship (MTD)

Cornell Lab of Ornithology Athena Fund (MTD)

Smithsonian Tropical Research Institute (MTD)

## Author contributions

Conceptualization: MSD, ANR, SDM

Methodology: MSD, MTD, ANR, VB, EB, SDM

Investigation: MTD, ANR, MSD, SDM

Data curation: EB, MTD, MSD, ANR

Formal analysis: EB, MTD, MSD, SDM, ANR

Visualization: MSD, SDM, VB, EB, MTD, ANR

Funding acquisition: ANR, MSD, MTD, VP

Project administration: MSD, ANR, VP

Supervision: ANR, MSD, VP

Writing – original draft: MSD, ANR, MTD, EB, SDM, VB

Writing – review & editing: ANR, MTD, MSD, EB, SDM, VB

## Competing interests

Authors declare that they have no competing interests

## Data and materials availability

Representative data on each species will be (at the time of final publication) available for open downloading at library.fisheyecollaborative.org for use under a CC BY-NC-ND license. The metadata for all media specimens follows Darwin Core standards and will be published on the Global Biodiversity Information Facility (GBIF). Additional data are available by request. All software algorithms are in the public domain and are referenced.

## Supplementary Information

According to an analysis by FishSounds.net, 1048 fish species have some published descriptions of acoustic behavior. Some of these published accounts include recordings, some have quantitative measures such as waveforms or spectrograms, and some are simply descriptive. In our review of fish sounds, we limited the scope of our search to reef-associated fish of the Caribbean by assessing only species with “reef” in the habitat type within the Smithsonian’s “Shorefishes of the Greater Caribbean” (Robertson and Van Tassell 2023) online information system. Combining the recordings in fishsounds.net with sounds we have found from other online sources (Vigness-Raposa et al. 2012, Schärer-Umpierre et al. 2019, Hatch et al. 2024), we found 174 species that have published acoustic behavior, of which only 132 have publicly available recordings of active sound production. Some of the documented sounds may have sound recordings held by researchers but not publicly available. There are likely recordings we could not find, reinforcing the need for organized databases for those seeking to use these recordings for their research (Parsons et al. 2022, Looby et al. 2023b).

We examined these existing sound recordings for Caribbean reef fish and categorized them by the degree to which they might reflect natural behaviors. Where multiple types of recordings were available for a species, we categorized them based on the recording made under the most natural circumstances and/or with the fewest stressors. Likewise, when notes for a recording indicated both free behaviors and unnatural stimuli, we categorized recordings as free behaviors. For the small number of fish for which sound recordings are freely available online (132), the majority (88) were recorded with electrical stimulation, manual stimulation (squeezing), or other handling that would result in great duress for the specimen. These techniques establish that the species has the ability to make sound, but the sound recordings are exceedingly unlikely to represent the type of natural behaviors that can be used to interpret a natural soundscape. A smaller fraction (11) of the recordings were obtained under mild duress, wherein fish were subjected to unnatural stressors resulting in annoyance, escape sounds, or territorial behaviors. This included stimuli such as prodding fish with a stick, inducing inflation in puffers, or introducing heterospecifics to experimental tanks. These recordings may reflect natural recordings, at least for fish in some natural peril. Some species (22) have had spontaneous sounds recorded in captivity. While these were typically recorded in tanks, some specimens were also kept in outdoor pens or bags suspended in water. These recordings likely reflect some natural behavior, but the behavioral context is not always specified. Generally, captive recordings are difficult to extrapolate to natural circumstances. The handling, enclosed space, human presence, and the recording environment may be stressors for the fish. For some species, natural behaviors are uncommon in captivity (Rice et al. 2022). Additionally, the differences in acoustics between glass tanks and the ocean may make tank recordings inaccurate representations of how fish sound in nature (Duncan et al. 2016).

Not all sounds produced by fish are vocalizations. Many references and recordings specify passive sound production from behaviors such as sudden movement or chewing. It is likely that most, if not all, species produce passive sounds during these behaviors. This does not make a species soniferous. We did not include recordings of movement or chewing in this analysis. It is unlikely that sounds from movement and chewing can be identified to a species level, however, a larger dataset of feeding sounds may reveal whether these recordings are useful for PAM analysis. Some more distinct feeding sounds, such as grazing sounds from parrotfish, may be identifiable to the family or genus level and are valuable indicators of herbivory on coral reefs. We found no such species-specific grazing sounds among previous recordings, so we have included grazing sounds in our library.

Of all recorded species, only a handful (11) have natural behaviors recorded in a natural environment. These recordings would be the best representations of spontaneous fish sounds as they do not face the drawbacks of direct stressors affecting fish behavior or aberrant tank acoustics. Most of these natural sounds are from fish in well-documented soniferous families, including Batrachoididae (toadfishes) and Serranidae (groupers). These recordings are the most useful for PAM analysis but are scarce. We aim to expand this number to create a more robust toolkit for using PAM on Caribbean reefs.

While captive recordings with or without additional stressors have drawbacks, they are not necessarily unusable for PAM analysis. Some of these sounds are likely similar to those produced in natural settings. Captive recordings may still be of interest to bioacousticians, especially those documenting free behaviors, as they can be used to corroborate new recordings of species-specific sounds. See Supplementary Table 1 for more information on previous recordings of spontaneous fish sounds in nature and captivity.

**Supplementary Table 1:**
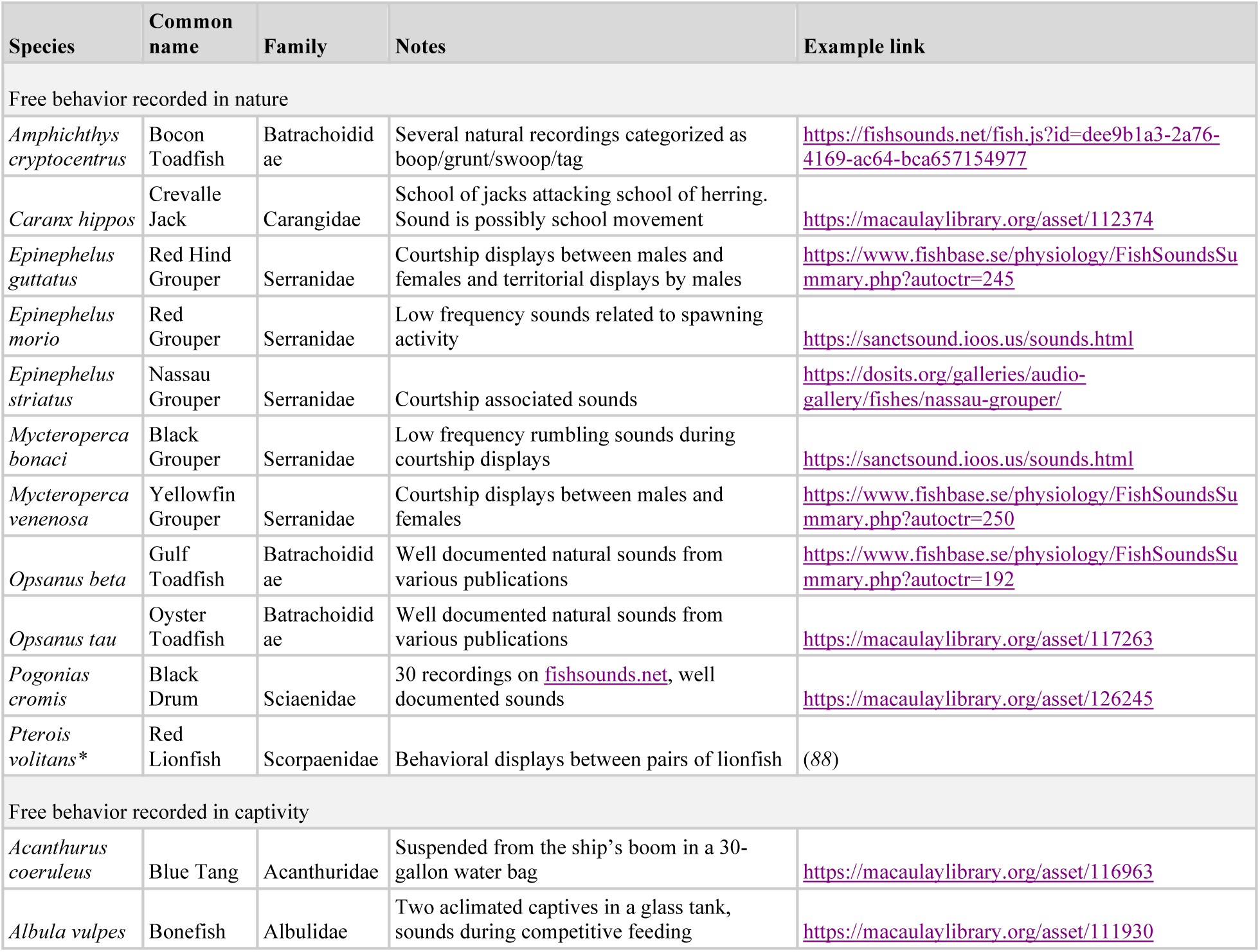
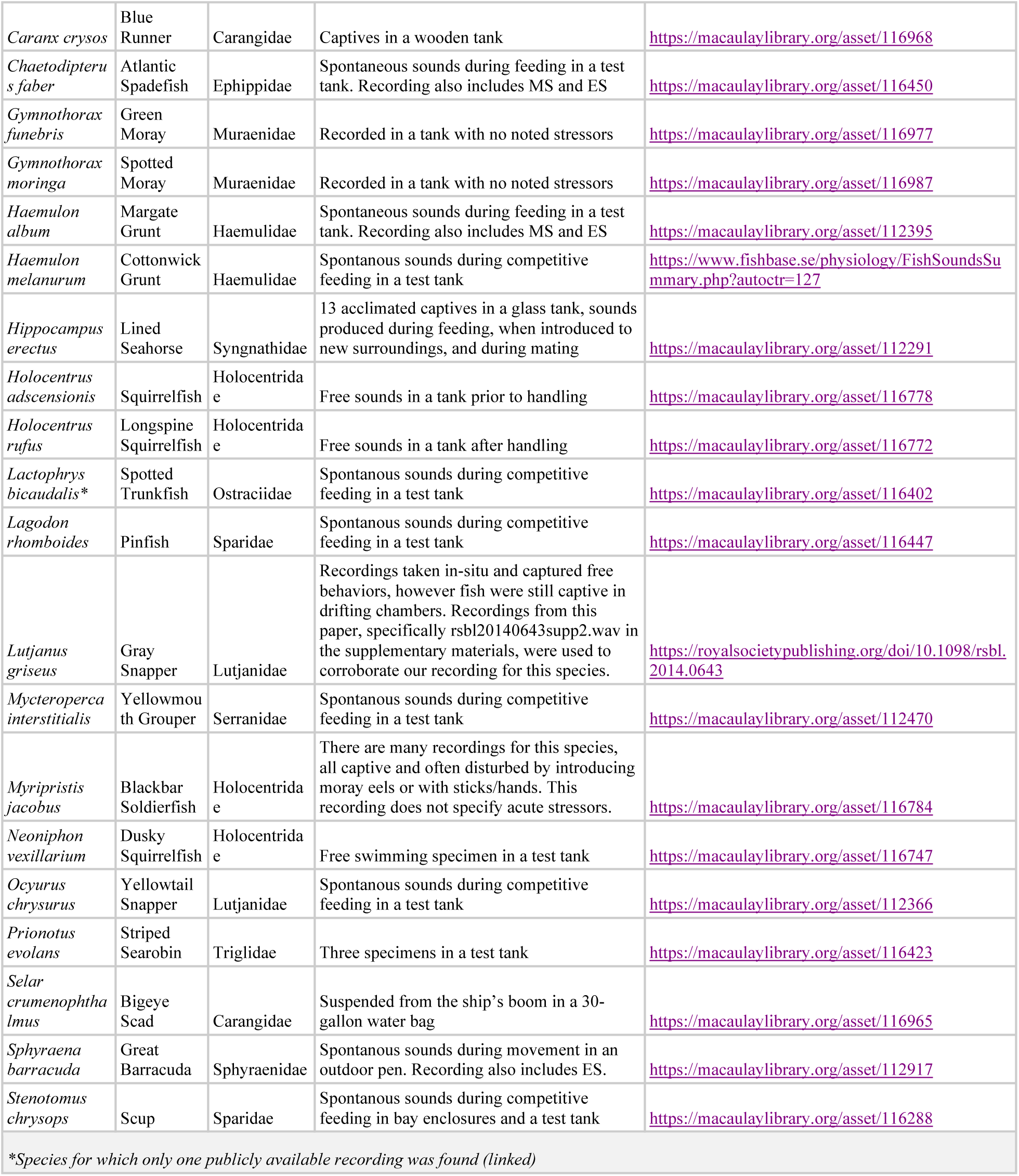
Summary of spontaneous fish sounds. These records are divided into free behaviors recorded in nature and those recorded in captivity. All of these species have been recorded making spontaneous sounds without additional stimuli. Some media notes described recordings as containing both spontaneous sounds and sounds elicited through duress (electric or manual stimulation). It is unclear which sections of the recordings are spontaneous. Most species have several recordings, sometimes from different sources available across public repositories (https://www.fishsounds.net, https://dosits.org/galleries/audio-gallery/fishes/, and https://sanctsound.ioos.us/sounds.html) and other publications, cited below. In these cases, we provided only a single example link to a sound representative of free behaviors; however, it should be noted that other recordings that may be beneficial for PAM analysis exist online.

## Supplementary Movie Captions

**Supplementary Movie 1: Ascribing sounds to individuals:** Video corresponding to Figure 5. Demonstrates our ability to ascribe sounds to individuals.

**Supplementary Movie 2. Ascribing sounds to species:** Video corresponding to Figure 6. Examples of sounds we have ascribed to species showing the complete detection events.

**Supplementary Movie 3. Determining the behavioral context of sounds.** Sounds and associated behaviors of (**A**) *Halichoeres garnoti*, Yellowhead Wrasse (**B**) *Stegastes partitus*, Bicolor Damselfish, (**C**) *Myripristis jacobus*, Blackbar Soldierfish, (**D**) *Sparisoma viride*, Stoplight Parrotfish, (**E**) *Acanthurus coeruleus*, Blue Tang, (**F**) *Aulostomus maculatus*, Atlantic Trumpetfish, and (**G**) *Scarus iseri*, Striped Parrotfish. The behavioral context is often clear, while others are still unknown.

## Notes

### Competing Interest Statement

The authors have declared no competing interest.

### Summary of Updates

Title updated, references updated, text copy-edited and updated. Supplemental files updated.

